# Cell-type targeted CRISPR/Cas9 Clock knockdown in mouse VTA dopamine neurons alter sleep, behavior, and cellular excitability

**DOI:** 10.64898/2026.06.03.730017

**Authors:** Yeon Ha Ju, Janet Y. Zhao, Setareh Sianati, Christine Egebjerg, Oscar C. González, Abby Criswell, Chaeyun Yuni Lee, Kaitlyn Du, Charlotte E. Luff, Joanna Stralka, Valentina Knapp, Luis de Lecea, Yevgenia Kozorovitskiy, Julie A. Kauer, Lief E. Fenno

## Abstract

Bipolar disorder (BD) is a severe psychiatric disease characterized by recurrent mania, depression, and circadian rhythm disruption. Among circadian regulators implicated in mood-related dysfunction, Clock has emerged as a particularly strong mechanistic candidate. However, cell type-specific functions of Clock within mood-relevant circuits remain incompletely defined. Here, we developed and applied a Cre-dependent AAV-SaCas9 gene-editing strategy to disrupt Clock selectively in ventral tegmental area dopamine neurons. We first established a rapid *in vitro* screening pipeline for guide RNA selection that accurately predicted *in vivo* editing efficiency. We then targeted Clock *in vivo* using a single AAV-based editing strategy and observed robust titer-dependent reduction of Clock expression, by targeted sequencing, *in situ* hybridization, and immunohistochemistry. We assessed the functional consequences of Clock disruption across analysis levels, including a behavioral battery, circadian and sleep-wake measurements using EEG and EMG, and electrophysiological recordings. These results establish a practical framework for rapid, cell-type-specific disruption of candidate psychiatric risk genes and provide a mechanistically grounded model for investigating how loss of Clock function in mesolimbic dopaminergic circuits contributes to BD-relevant phenotypes.

## Introduction

Bipolar disorder (BD) is a severe, polygenic psychiatric disease characterized by recurrent episodes of mania and depression. In addition to affective symptoms, disturbances in sleep and circadian timing are among the core features of BD^1–4^. Individuals living with BD commonly show altered circadian phase relationships, shortened or fragmented sleep, and an instability in daily activity rhythms^5–8^. Importantly, circadian and sleep disruptions frequently precede mood episodes and acute sleep deprivation can precipitate mania^9–11^, suggesting that circadian dysfunction may be a proximal contributor to mood destabilization rather than a mere downstream consequence of affective states and transitions^4,12^.

Circadian rhythm is governed by conserved transcriptional and translational feedback loops, where the core clock components, including Clock, Bmal1, Per, and Cry, generate near 24-hour oscillations in gene expression that coordinate physiological and behavioral rhythms across tissues^13–16^. Among the core circadian regulators, Clock has emerged as a particularly compelling candidate for mood-related dysfunction^17–19^. In preclinical models, the dominant-negative ClockΔ19 mutation produces a robust mania-like behavioral profile^20–23^, and subsequent work has shown that reducing Clock expression in the ventral tegmental area (VTA) is sufficient to induce marked alterations in locomotor, affective, and circadian phenotypes^20,22,24^. Together, these findings indicate that Clock is not simply correlated with BD-relevant behavior, but can causally influence neural systems that regulate reward, arousal, and mood states.

At the same time, an important mechanistic question remains unresolved. The VTA contains heterogeneous neuronal populations, including dopaminergic (DA), GABAergic, and glutamatergic cells^25,26^. Prior region-targeted manipulations do not establish whether loss of Clock specifically in VTA dopaminergic neurons is sufficient to drive these phenotypes^20,24^. Resolving this issue is important for linking circadian gene dysfunction to a specific neural substrate within the broad mesolimbic system. We hypothesized that selective disruption of Clock in VTA DA neurons would be sufficient to produce BD-relevant behavioral, circadian, and electrophysiological abnormalities. To test this hypothesis with cell-type resolution, we used Cre-dependent adeno-associated virus (AAV)-mediated CRISPR/SaCas9 to selectively disrupt Clock in VTA DA neurons. Targeted Clock loss drove hyperexcited behavioral, circadian, and electrophysiological states that parallel core features of human mania. By focusing on a single, well supported circadian regulator, this study establishes a conservative and mechanistically grounded model for assaying the sufficiency of BD risk gene disruption within restricted cell-types and neural circuits, to cause BD-relevant phenotypes.

## Results

### A single-AAV Cre-dependent SaCas9 system enables conditional gene editing and predicts *in vivo* sgRNA performance

To enable cell-type-specific genetic manipulation of BD-associated risk genes *in vivo*, we applied a previously described Cre-dependent SaCas9 system consisting of an inverted SaCas9 flanked by a double-floxed Inverted Orientation (DIO) switch^27,28^, alongside a U6-driven sgRNA cassette (AAV-CMV-DIO-SaCas9-U6-sgRNA; **Fig. 1A**)^29^. This single-vector configuration encodes a 5.1 kb transgene that exceeds the conventional 4.7 kb AAV packaging limit^30,31^, a property we return to when considering titer requirements for efficient *in vivo* editing.

**Figure 1.**
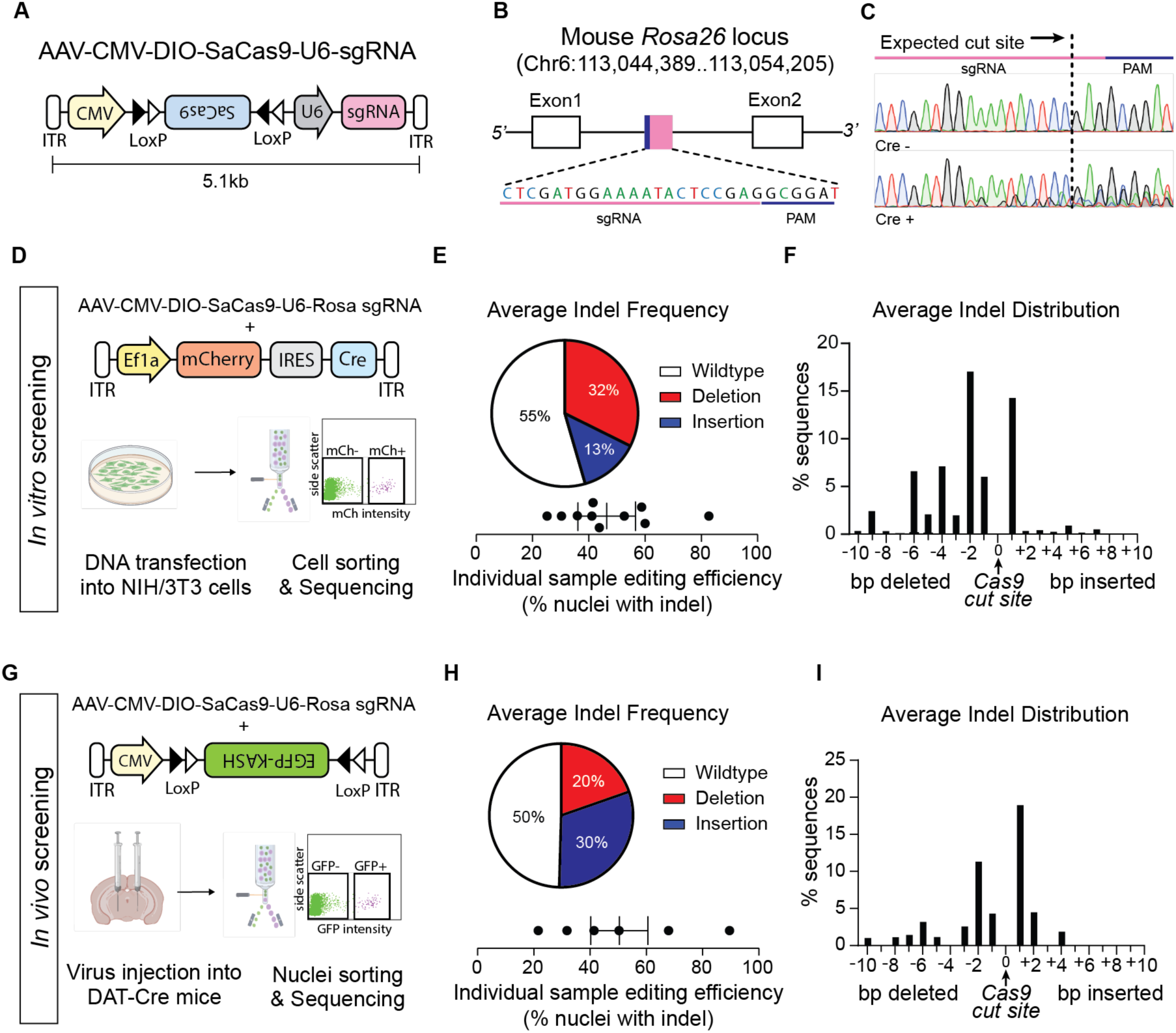
(**A**) Schematic of the Cre-dependent AAV-CMV-DIO-SaCas9-U6-sgRNA vector. (**B**) sgRNA target site within the Rosa26 locus, showing the sgRNA target sequence (red) and PAM sequence (blue); sgRNA target sequence is anti-sense relative to the gene coding sequence. (**C**) Representative sanger sequencing traces from Cre- and Cre+ samples, with the expected SaCas9 cut site indicated (arrow). (**D**) *In vitro* validation workflow. NIH/3T3 cells were co-transfected with AAV-CMV-DIO-SaCas9-U6-Rosa26 sgRNA and AAV-Ef1a-mCherry-IRES-Cre. mCherry-positive cells isolated by FACS and analyzed by amplicon sequencing. (**E**) Pie chart showing average indel frequency (wild-type, deletion, insertion), with individual sample editing efficiencies shown below. (**F**) Bar graph showing average distribution of indel sizes, grouped by deletion and insertion length in base pairs. (**G**) *In vivo* validation workflow. DAT-Cre mice were co-injected with AAV-DIO-KASH-EGFP and AAV-DIO-SaCas9-U6-Rosa26 sgRNA. (**H**) Pie chart showing the average indel frequency, with individual sample editing efficiencies shown below. (**I**) Bar graph showing average distribution of indel sizes, grouped by deletion and insertions (bp).

We first validated the system at the mouse Rosa26 locus, a well-characterized safe-harbor site commonly used for genome editing validation (**Fig. 1B**)^32,33^. We used a previously characterized SaCas9 sgRNA sequence^29^ that targets the intronic region between exon 1 and exon 2. We co-transfected AAV-CMV-DIO-SaCas9-U6-Rosa26 sgRNA vector with a Cre-expressing plasmid (pAAV-Ef1a-mCherry-IRES-Cre) into NIH/3T3 mouse cells. Three days post-transfection, mCherry+ cells were isolated by fluorescence-activated cell sorting (FACS) to restrict analysis to Cre-expressing cells (**Supplemental Fig. 1A**). Sanger sequencing of PCR amplicons from transfected NIH/3T3 cells confirmed indel formation at the expected cut site approximately 3 bp upstream of the PAM sequence exclusively in the presence of Cre recombinase, consistent with known SaCas9 cleavage properties (**Fig. 1C,D**)^34^, validating Cre-dependent activation and function of the system. Amplicon sequencing of the Rosa26 locus revealed an average indel frequency of 45%, comprising 32% deletions and 13% insertions (**Fig. 1E, F**).

Next, we assessed *in vivo* editing by stereotactic injection of AAV-CMV-DIO-SaCas9-Rosa26 sgRNA co-delivered with AAV-CAG-DIO-EGFP-KASH into the VTA of DAT-Cre mice (**Fig. 1G**). Four weeks post-injection, single nuclei were isolated and GFP+ nuclei collected by FACS, restricting genomic analysis to Cre-expressing dopaminergic neurons. Genomic sequencing revealed an average indel frequency of approximately 50%, closely mirroring *in vitro* observations (**Fig. 1H, I, J**).

Taken together, these data replicate previously published results^29^ targeting the Rosa26 locus using Cre-dependent AAV-SaCas9, and provide preliminary data suggesting that our approach to selecting sgRNA candidates enabled useful concordance between *in vitro* and *in vivo* editing efficiencies. Having identified a platform approach for targeted gene disruption, we next turned our attention to the BD-associated gene Clock.

### Targeting Clock in VTA dopamine neurons validates *in vivo* editing and reveals titer dependence

Genetic and biological evidence link Clock, circadian dysfunction, and BD^19,35,36^, while prior studies have illustrated that Clock disruption is sufficient to induce mania-like phenotypes in rodents^20,24^. To disrupt Clock using CRISPR, we designed sgRNA sequences by aligning all NCBI-documented splice variants and prioritizing candidates based on efficiency scores and minimal predicted off-target activity (CRISPOR, CHOPCHOP)^37–41^, generating three candidate sgRNAs targeting distinct exons encoding functionally critical protein domains: sgRNA #1 (Exon 9), sgRNA #2 (Exon 10), and sgRNA #3 (Exon 16) (**Fig. 2A**). *In vivo* editing efficiency in the VTA of DAT-Cre mice was assessed as described above. sgRNA #1 and sgRNA #2 yielded average indel frequencies of 39% and 40%, respectively. sgRNA #3 emerged as the superior candidate, achieving an average indel frequency of 69%, with a predominant deletion rate of 63% (**Fig 2B-D**). These results identified sgRNA #3 as the lead Clock sgRNA for subsequent experiments. For the subset of Clock sgRNAs also evaluated *in vitro*, we observed the same relative pattern seen *in vivo*, with sgRNA #3 showing higher editing efficiency than sgRNA #2 (44.6% vs. 28.8%, respectively; **Supplementary Fig. 1B**). To further assess whether this relationship extended across guides more broadly, we compared matched *in vitro* and *in vivo* editing efficiencies for the subset of sgRNAs tested in both contexts and observed a positive correlation between the two measures (R² = 0.4886, p = 0.0037; **Supplementary Fig. 1C**). Together, these findings suggest that *in vitro* screening can serve as a practical approach for prioritizing sgRNAs with stronger *in vivo* editing performance.

**Figure 2.**
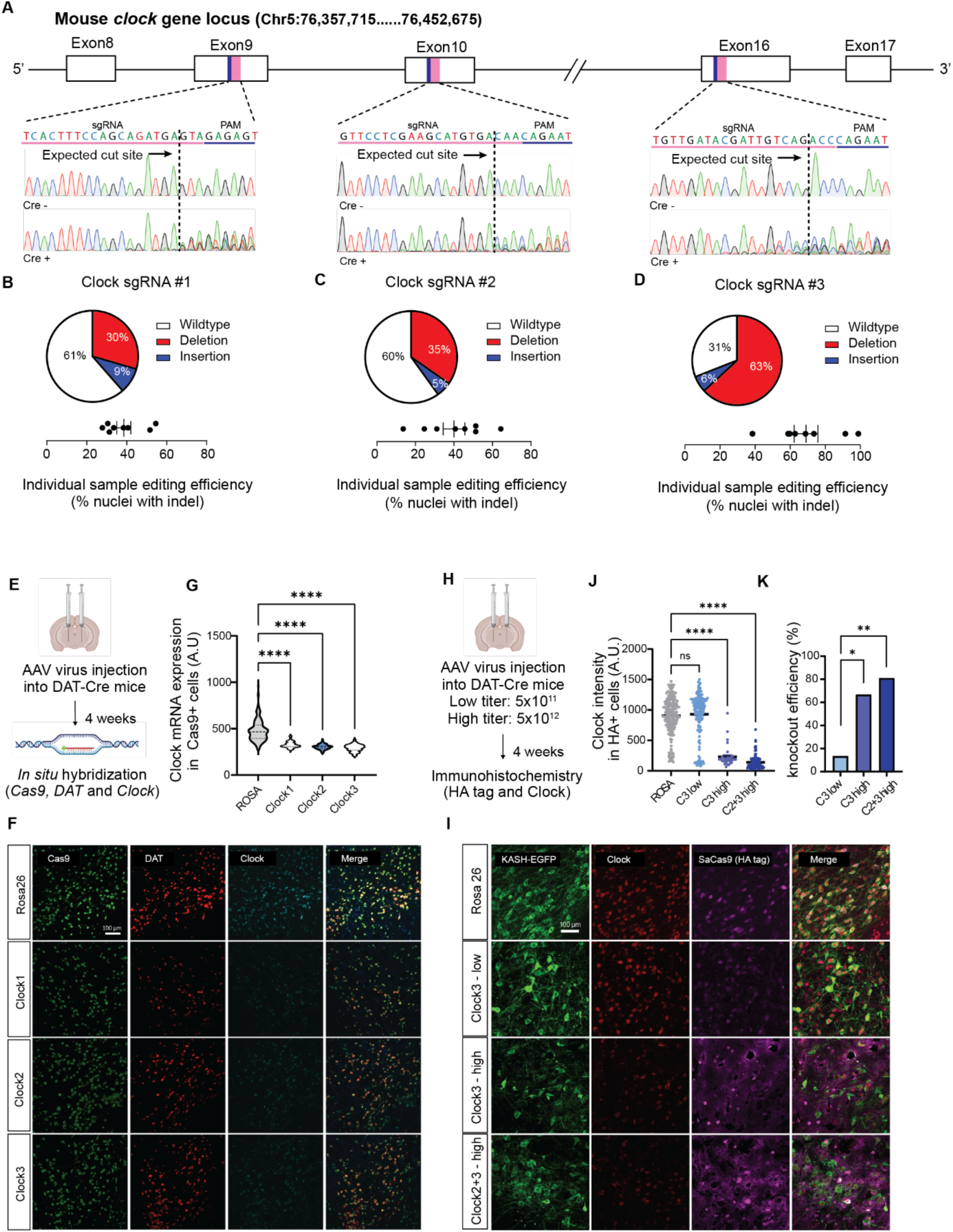
**(A)** Mouse *clock* gene structure with sgRNA target sites in Exons 9, 10, and 16; sgRNA (blue) and PAM (red) sequences indicated. sgRNA target sequence is anti-sense relative to the gene coding sequence. Sanger traces from GFP+ nuclei confirm editing at each site; GFP- traces not shown. (**B-D**) Indel distributions for Clock sgRNAs #1-#3 from *in vivo* virally-transduced nuclei as pie charts and scatter plots of individual editing efficiencies. sgRNA #3 achieved 69% editing efficiency. (**E**) *In situ* hybridization quantifying Clock mRNA in Cas9-expressing VTA neurons. (**F**) Representative fluorescence images for Cas9 (green), DAT (red) and Clock (cyan). Scale bar = 100 μm. (**G**) Violin plot quantifying Clock mRNA levels in Cas9-expressing cells. Clock-targeted sgRNA significantly reduced mRNA relative to Rosa26 control. (**H**) Experimental design for immunohistochemistry (IHC). Virus expressing Cas9 and sgRNAs targeting Clock (C3 or C2+3) was injected at different titers (low: 5 × 10¹¹, high: 5 × 10¹²). After 4 weeks, brain sections were processed for IHC. **(I**) Representative IHC images showing Clock (red), and HA-tag (Cas9+, magenta). Scale bar = 100 μm. (**J**) Clock intensity in HA+ (Cas9+) cells. (**K**) Bar graph showing knockdown efficiency. *p< 0.05, **p<0.01, ****p<0.0001

To confirm that the genomic indels translated into depletion of Clock gene products, we performed independent histological and molecular validation in the VTA of DAT-Cre mice 4 weeks after AAV injection. *In situ* hybridization using probes against *Clock*, *Cas9*, and *DAT* revealed significant attenuation of Clock mRNA in Cas9-positive cells for all three sgRNA candidates relative to the Rosa26 control, confirming robust reduction of Clock transcript at the single-cell level (**Fig. 2E-G**).

We next asked whether the observed transcript-level disruption translated into protein loss, and whether editing efficiency was sensitive to viral titer, a question motivated by the supra-threshold payload size of the single-vector system. Immunohistochemistry using antibodies against Clock and the HA-tagged SaCas9 was performed following injection of sgRNA #3 at low titer (5×10^11^ vg/mL), high titer (5×10^12^ vg/mL), or after co-injection of independent single-guide viruses targeting sgRNA #2 and #3 at high titer (**Fig. 2H, I**). Low-titer injection did not significantly reduce Clock protein intensity compared to the Rosa26 controls. High-titer delivery of sgRNA #3 produced a significant reduction in Clock protein levels, and co-injection of two independent sgRNA #2 and #3 viruses at high titer further enhanced knockdown, achieving approximately 80% knockdown efficiency (**Fig. 2J, K**). Together, these data show that effective protein-level disruption in this single-vector system is strongly dependent on viral titer and can be further improved by combining independent guides targeting distinct exons.

### Exome sequencing detects no evidence of off-target editing by Clock or Rosa26 sgRNAs in vivo

To assess potential CRISPR/SaCas9 off-target editing *in vivo*, we performed exome sequencing on genomic DNA isolated from sorted cells obtained from mice injected with each of the four sgRNA conditions, Clock1, Clock2, Clock3, and Rosa26. For each condition, GFP-positive nuclei, indicating viral transduction and Cre-dependent gene expression were collected from 4 mice. GFP-negative nuclei isolated from the same animals served as within-animal wild-type controls and were processed for exome sequencing. Across all 16 mice and five analyzed groups, exome sequencing identified 1,306 unique indels relative to the mouse reference genome (GRCm38/mm10). When counted within each condition, 443 indels were detected in Rosa26 sgRNA samples, 315 in Clock1 sgRNA samples, 430 in Clock2 sgRNA samples, 349 in Clock3 sgRNA samples, and 837 in wild-type samples; the higher raw count in wild-type samples reflects the larger aggregated sample size of wild-type nuclei (n = 16 mice, pooled across all four cohorts) compared to each individual sgRNA condition (n = 4 mice each). Of the 1,306 total indels, 152 were shared across all five groups, including wild-type cells, and 268 were present in at least two groups (**Table 1**). These shared indels were excluded from further analysis, as were 16 indels annotated as common variants. After filtering, 445 indels were unique to wild-type samples, whereas 150, 59, 151, and 81 indels were unique to the Rosa26, Clock1, Clock2, and Clock3 sgRNA conditions, respectively (**Table 2**).

**Table 1.** Summary of identified potential off-target sites.

| Mismatches | ROSA | Clock1 | Clock2 | Clock3 |
| --- | --- | --- | --- | --- |
| 0 | 3 | 5 | 1 | 1 |
| 1 | 4 | 4 | 7 | 5 |
| 2 | 9 | 8 | 10 | 18 |
| 3 | 51 | 89 | 117 | 345 |
| 4 | 910 | 1414 | 1649 | 2314 |
| 5 | 3503 | 3951 | 4063 | 4123 |
| 6 | 4295 | 4452 | 4577 | 5000 |
| 7 | 5000 | 5000 | 5000 | 5000 |

**Table 2.** Summary of detected indels.

| Conditions | Indels |
| --- | --- |
| All | 152 |
| 2+ | 268 |
| ROSA only | 150 |
| Clock1 only | 59 |
| Clock2 only | 151 |
| Clock3 only | 81 |
| WT only | 445 |
| Total | 1306 |

To identify candidate off-target sites, we used Cas-OFFinder to predict genomic loci with sequence similarity to each sgRNA target^42,43^. Search was performed allowing up to seven mismatches and up to 2 bp DNA or RNA bulges, corresponding to the maximum mismatch and bulge settings supported by the algorithm. This analysis identified 13,775, 14,923, 15,424, and 16,806 potential off-target sites for Rosa26, Clock1, Clock2, and Clock3 sgRNAs, respectively. We then compared these predicted sites with the filtered indels detected by exome sequencing. On-target editing was detected in samples from the Clock1, Clock2, and Clock3 conditions. In contrast, the Rosa26 on-target site was not evaluable by exome sequencing because it lies within an intronic region not captured by the exome panel. Importantly, none of the filtered indels overlapped predicted off-target sites for any sgRNA. Together, these data provide no evidence of exome-detectable off-target editing under the conditions tested.

### Clock disruption in VTA dopamine neurons produces BD-relevant behavioral phenotypes and alters sleep architecture

We next asked whether selective, early postnatal Clock disruption in VTA dopaminergic neurons is sufficient to drive BD-relevant behavioral abnormalities *in vivo*. Prior studies have shown that Clock manipulation in the VTA of adult animals can induce mania-like phenotypes^20,24^, but the circuit-specific contribution of Clock loss within DA neurons remains less clear. To examine the contribution of Clock knockdown starting in early postnatal development, we performed neonatal bilateral AAV injections targeting the VTA in DAT-Cre mice, using either control (Rosa26) or Clock3 sgRNA CRISPR AAVs. Each was co-delivered with a Cre-dependent fluorescent reporter AAV expressing tdTomato. After 8 weeks of expression, mice underwent behavioral testing to monitor rodent analogs of mania- and depression-like spectrum behaviors, or EEG/EMG recordings to assess sleep architecture. Injection targeting and viral expression were validated post hoc for all mice as illustrated in **Supplementary Fig. 2**.

To assess general motor and anxiety-related behavior, we performed an open field test. At baseline, mice with Clock CRISPR viruses in VTA DA neurons displayed reduced anxiety-like behavior in an open-field locomotion assay, evidenced by a significant increase in time spent in the center of the arena (**Fig. 3A**). This loss of species-typical thigmotaxis was not driven by increased locomotion, as mobile speed and average acceleration did not differ significantly between Rosa26 controls and Clock KD mice (**Fig. 3A**). Consistent with prior reports of VTA Clock manipulation^20,24^ and building upon them, these findings indicate that Clock disruption in VTA dopaminergic neurons alone is sufficient to recapitulate an anti-anxiety-like phenotype, without increasing baseline locomotion.

**Figure 3.**
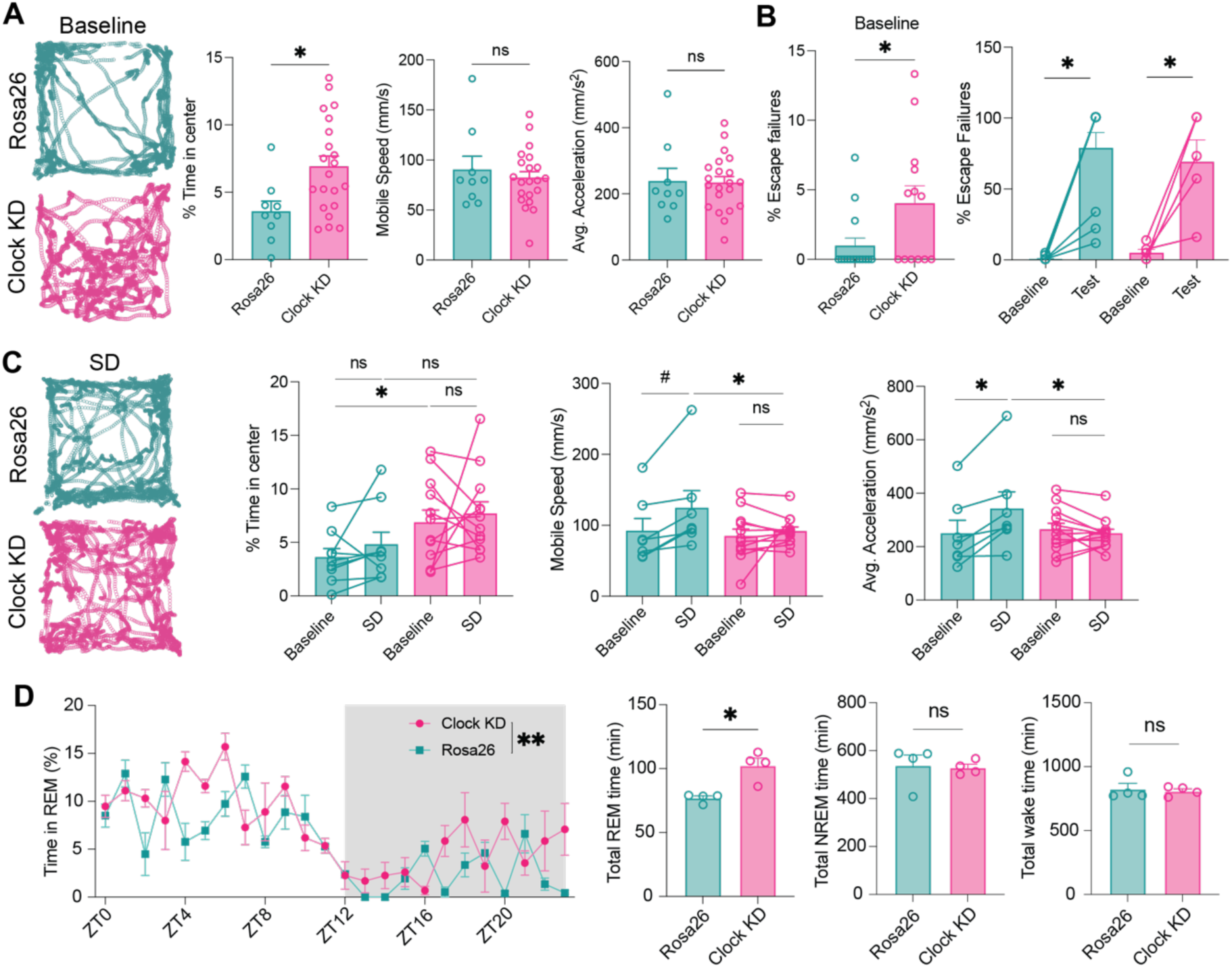
(**A**) Open field locomotor behavior. Example trajectories over 5 min (left) and quantified time in center, mobile speed, and average acceleration for Rosa26 (n=9) and Clock (n=21) animals (right). Clock KD animals spend significantly more time in the center, consistent with reduced anxiety-like behavior. Mobile speed and acceleration do not differ. (**B**) Escape failures in a learned helplessness paradigm are elevated in Clock KD mice (n=12) relative to Rosa26 controls (n=9) at baseline but not after 2 induction sessions. (**C**) Open field behavior after 12 hours of sleep deprivation (SD). Rosa26 controls (n=9) show significant increases in average acceleration following SD; Clock KD mice (n=12) show no such changes, revealing a selective failure of normal SD-associated behavioral responses. Time in the center does not increase after SD. (**D**) Sleep architecture measured by EEG/EMG recording over 24 hours. Clock KD mice (n=4) have a significant increase in total REM sleep duration relative to Rosa26 controls (n=4). Total NREM and wake duration are not significantly affected. ns = not significant, #p< 0.1, *p< 0.05, **p<0.01.

Next, we assessed depression spectrum-relevant behavior using a classical learned helplessness paradigm. Prior to an aversive learning experience, wild-type mice escape nearly all foot shocks under baseline conditions^44,45^. Here, in contrast, mice with disrupted Clock in VTA DA neurons showed a greater number of escape failures on first exposure to the task with no change in escape failures following induction sessions (**Fig. 3B**). These data, consistent with decreased behavioral resilience, indicate that selective Clock disruption in VTA dopaminergic circuitry is sufficient to produce coexisting behavioral abnormalities across anxiety-related and stress-coping domains.

To further probe the impact of Clock knockdown on experience-driven adaptations, we next assessed how mice respond to sleep disruption in the absence of intact Clock function in VTA DA neurons. A subset of subjects underwent 12 hours of sleep deprivation (SD), while the remaining cohort remained in their homecage (**Supplementary Fig. 3A**). Consistent with our prior work^45^, Rosa26 control mice showed increased locomotion following SD (**Fig. 3C**) that was not seen in mice who were allowed to remain in their homecage overnight **(Supplementary Fig. 3B**). In contrast, mice with Clock disruption in VTA dopaminergic neurons did not exhibit this SD-induced behavioral change (**Fig. 3C**) and displayed similar locomotor behaviors to Clock KD homecage controls (**Supplementary Fig. 3B**). In the learned helplessness paradigm, there was no change in the number of escape failures in Clock KD mice following 12 hours of SD (**Supplementary Fig. 3B**), in contrast to what has been shown previously in wild-type mice^45^. These data suggest that loss of Clock in VTA dopaminergic neurons alters sensitivity to SD-associated transient adaptive behavioral responses.

We next asked whether early postnatal Clock disruption in VTA dopaminergic neurons impacts sleep architecture. EEG and EMG recordings collected across 24 hours revealed that Clock KD mice spent significantly more time in REM sleep than Rosa26 controls (mixed ANOVA, p < 0.01) (**Fig. 3D**). Total REM duration was increased, whereas total NREM sleep and wake time were unchanged. In contrast, when Clock disruption was induced in adult mice, 6-8 weeks of age, no significant changes in sleep architecture were detected (**Supplementary Fig. 3C**). Together, these results indicate that Clock disruption in VTA dopaminergic neurons is sufficient to alter BD-relevant behavioral domains and REM sleep architecture, with the effects on sleep structure dependent on the age when Clock disruption occurred.

### Clock disruption increases VTA DA neuron excitability and alters responses to sleep disruption

To determine how Clock disruption affects the intrinsic excitability of VTA dopaminergic neurons, we performed whole-cell recordings from fluorophore-expressing DA neurons in acute midbrain slices prepared from Clock3 sgRNA CRISPR KD mice. Clock-disrupted neurons exhibited increased excitability, characterized by a depolarized resting membrane potential and increased spike output in response to depolarizing current injections, indicating enhanced responsiveness to excitatory input (**Fig. 4A-D**). Notably, acute sleep deprivation in control (Rosa26) mice produced a similar increase in DA neuron excitability, paralleling the effects observed following Clock knockdown, while SD in mice with Clock3 sgRNA CRISPR KD did not further increase excitability (**Fig. 4C-D**). In contrast, all other measured intrinsic electrophysiological properties were unchanged relative to Rosa26 controls, including input resistance, action potential threshold, and H-current (**Fig. 4D)**.

**Figure 4.**
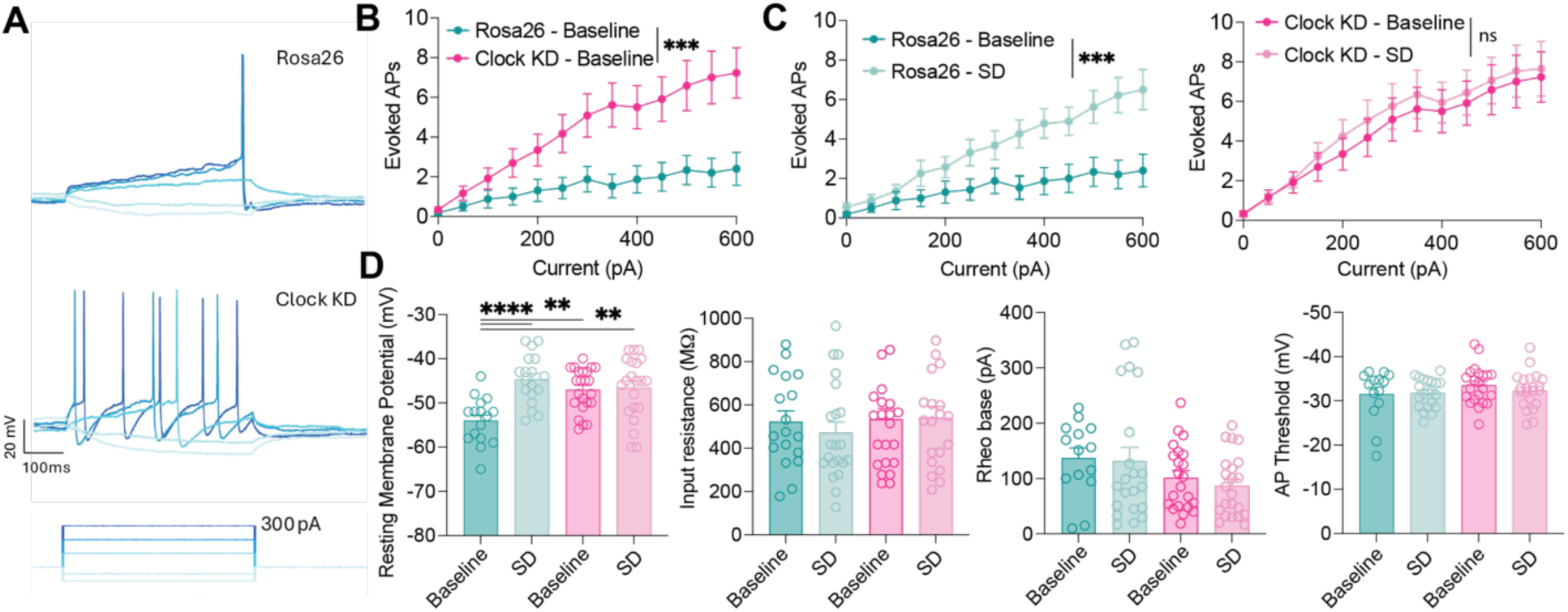
(**A**) Representative whole-cell patch-clamp traces from VTA DA neurons in acute brain slices, showing subthreshold and suprathreshold responses to current injection in Rosa26 (top) and Clock (bottom) knockdown. CRISPR transfected DA neurons were identified by fluorescence from a co-injected Cre-dependent tdTomato reporter virus. (**B**) Clock knockdown (KD) neurons (n=16) fire significantly more evoked action potential than Rosa26 controls (n=23) across injected current steps. (**C**) The same non-sleep-deprived Rosa26 and Clock KD datasets shown in **B** are replotted alongside sleep-deprived groups (Rosa26 SD, n=20; Clock KD SD, n=22). Sleep deprivation significantly increases evoked firing in Rosa26 controls but did not further increase firing in Clock KD neurons suggesting a selective failure of homeostatic excitability upregulation in Clock KD animals. (**D**) Resting membrane potential is significantly depolarized in Clock KD relative to Rosa26. SD depolarizes resting membrane potential in Rosa26 controls, but in CLOCK KD neurons is unchanged by SD. Input resistance, rheobase, and AP threshold do not differ across groups. **p<0.01, ***p<0.001, ****p<0.0001.

Next, to evaluate whether the observed behavioral and electrophysiological phenotypes were accompanied by changes in HPA axis activity^46^ we measured serum corticosterone at ZT12, the beginning of the active phase, in control and sleep-deprived Rosa26 and Clock KD animals. Corticosterone levels did not differ significantly across groups (**Supplementary Fig. 3D**), suggesting that Clock disruption in VTA dopaminergic neurons using this approach is not accompanied by detectable changes in basal circulating corticosterone at the start of the active phase.

## Discussion

Here, we applied a Cre-dependent AAV-SaCas9 strategy to disrupt Clock selectively in VTA dopaminergic neurons and found convergent effects across behavior, sleep-related phenotypes, and intrinsic neuronal excitability. These findings support the conclusion that Clock function in VTA dopamine neurons contributes to the regulation of affective state, arousal-related processes, and neuronal excitability. More broadly, the results identify a cell type-specific circadian mechanism within the mesolimbic dopamine system that is sufficient to generate phenotypes broadly relevant to BD.

An important implication of this study is that the effects of Clock disruption can be localized more precisely than in prior global or region-wide models. Previous work has established that altered Clock function influences mood-related behavior and that VTA-directed manipulations are sufficient to produce prominent behavioral abnormalities^20,24^. Our results extend that framework by showing that somatic disruption of endogenous Clock specifically in VTA dopaminergic neurons is sufficient to alter circuit function and behavioral output. This narrows the relevant cellular and neural circuit substrate through which circadian gene dysfunction may influence mood-related states.

At the physiological level, Clock disruption increased the intrinsic excitability of VTA dopaminergic neurons, supporting the idea that circadian molecular programs constrain dopaminergic activity in the adult brain^21,35^. Because VTA dopamine neurons regulate reward, motivation, and arousal, a shift toward elevated excitability may provide a mechanistic basis for the altered behavioral state observed after Clock disruption^47,48^. The observation that acute sleep deprivation produced partially overlapping electrophysiological effects in control neurons further suggests that loss of Clock may mimic one component of a sleep-disrupted circuit state, rather than globally reproducing all consequences of sleep loss.

At the behavioral level, the observed phenotypes did not map cleanly onto a single mania-like or depression-like category. Instead, Clock disruption produced changes across multiple domains, including reduced anxiety-like behavior, impaired stress coping, and altered responses to sleep deprivation. This pattern is more consistent with dysregulation of behavioral state than with unitary behavioral syndrome. Indeed, mixed states have been reported in a subset of individuals living with BD^49,50^. Accordingly, we interpret this model not as a full recapitulation of BD, but as a mechanistically defined perturbation that captures one circuit-level component of BD-relevant pathology.

Sleep phenotypes further refined this interpretation. We observed a selective increase in REM sleep without broad alterations in total wake or NREM time, suggesting that Clock function in VTA dopaminergic neurons influences specific components of sleep architecture rather than globally controlling sleep-wake organization. Based on the observation that VTA DA neurons are active during wakefulness and REM sleep^47^, the observed phenotype may reflect interactions of DA release with REM-generating neurons in the brainstem, or, alternatively, with sleep-modulating circuitry in the lateral hypothalamus^51^. Given the established links between REM regulation, emotional processing, and mood disorders^19,52^, this selective effect may represent one pathway through which circadian disruption alters behavioral state. At the same time, the relative specificity of the sleep phenotype indicates that disruption of Clock in adult VTA dopamine neurons alone is unlikely to account for the broader sleep and circadian abnormalities associated with BD, and that additional cell types, circuits, or developmental mechanisms are likely involved. The dependence of sleep phenotype on the developmental age of the subject at the time of Clock knockdown provides additional nuance, suggesting that both neurodevelopmental as well as homeostatic mechanisms may be impacted by aberrant gene function.

Several limitations should be considered. First, BD is highly polygenic^53,54^, and the disruption of Clock alone cannot capture the full genetic architecture of the disease. Second, CRISPR-mediated editing is expected to be mosaic, and variability in editing efficiency across neurons may contribute to phenotypic variability^55,56^. Third, while neonatal viral transduction captures the BD-relevant early developmental time window for viral expression, precise localization of transduction is more challenging in the neonates, especially when using Cre recombinase lines with broad expression. Finally, because our manipulation was restricted to one gene and a single cell population, the present model should be viewed as a reductionist test of a defined mechanistic hypothesis rather than a comprehensive disease model. Future studies will be needed to determine whether combinatorial perturbation of multiple BD-associated genes within shared neural circuits produces emergent phenotypes beyond those observed after single-gene disruptions and should further define how altered dopaminergic activity propagates through downstream targets to shape neural activity patterns and behavioral outcomes.

## Supplemental Figures

**Supplementary Figure 1.**
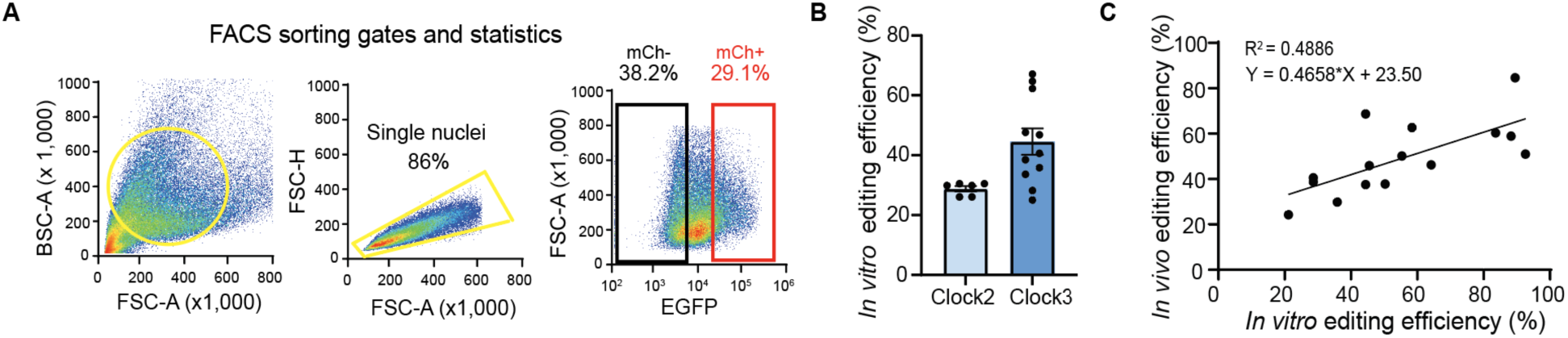
(**A**) Representative FACS gating strategy and sorting statistics for NIH3T3 cells. (**B**) *In vitro* editing efficiency for Clock sgRNA #2 and #3. Mean editing efficiencies were 28.8% for sgRNA #2 and 44.6% for sgRNA #3. (**C**) Correlation between *in vivo* and *in vitro* editing efficiencies across sgRNA candidates. Each dot represents one sgRNA candidate. Linear regression and coefficient of determination are shown (R^2^ = 0.4886, P = 0.0037, y=0.4658x+23.50).

**Supplementary Figure 2.**
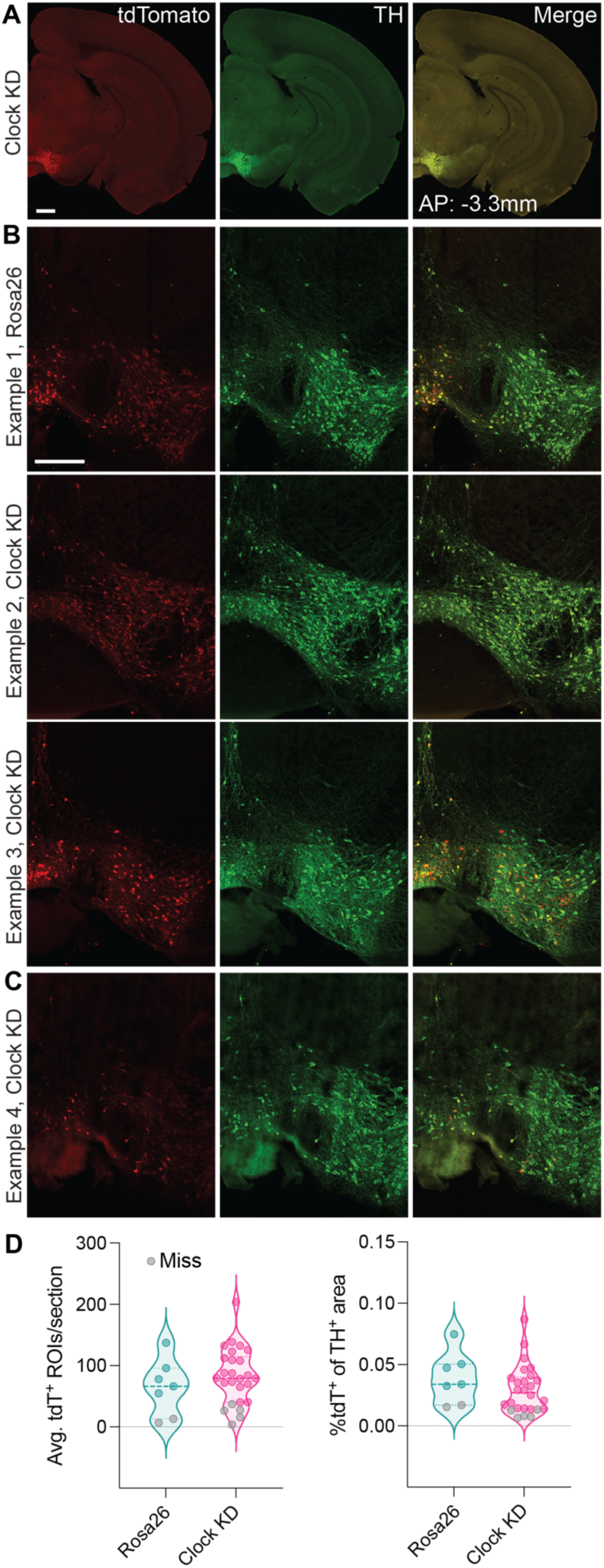
(**A**) tdTomato and tyrosine hydroxylase (TH) expression in a neonatally transduced Clock KD mouse. Scale bar, 500 μm. (**B**) Same as **A**, but example 1 is of a control animal and examples 2-3 are Clock KD mice. Scale bar, 250 μm. (**C**) tdTomato and TH expression of a Clock KD animal considered a missed injection and therefore not included in the behavior dataset. (**D**) Quantification of the number of tdT+ regions of interest (ROIs) per 60 μm section of the VTA (left). 2-3 sections were imaged per mouse and the number of tdT+ ROIs was averaged per section. Quantification of the percent of TH+ area that is also tdT+ for a single VTA section (right). Grey circles, using the threshold of the bottom quartile in the quantification on the left, are considered missed injections and not included in the behavior data set. Rosa26: n = 7; Clock KD: n = 26. No statistically significant difference between groups.

**Supplementary Figure 3.**
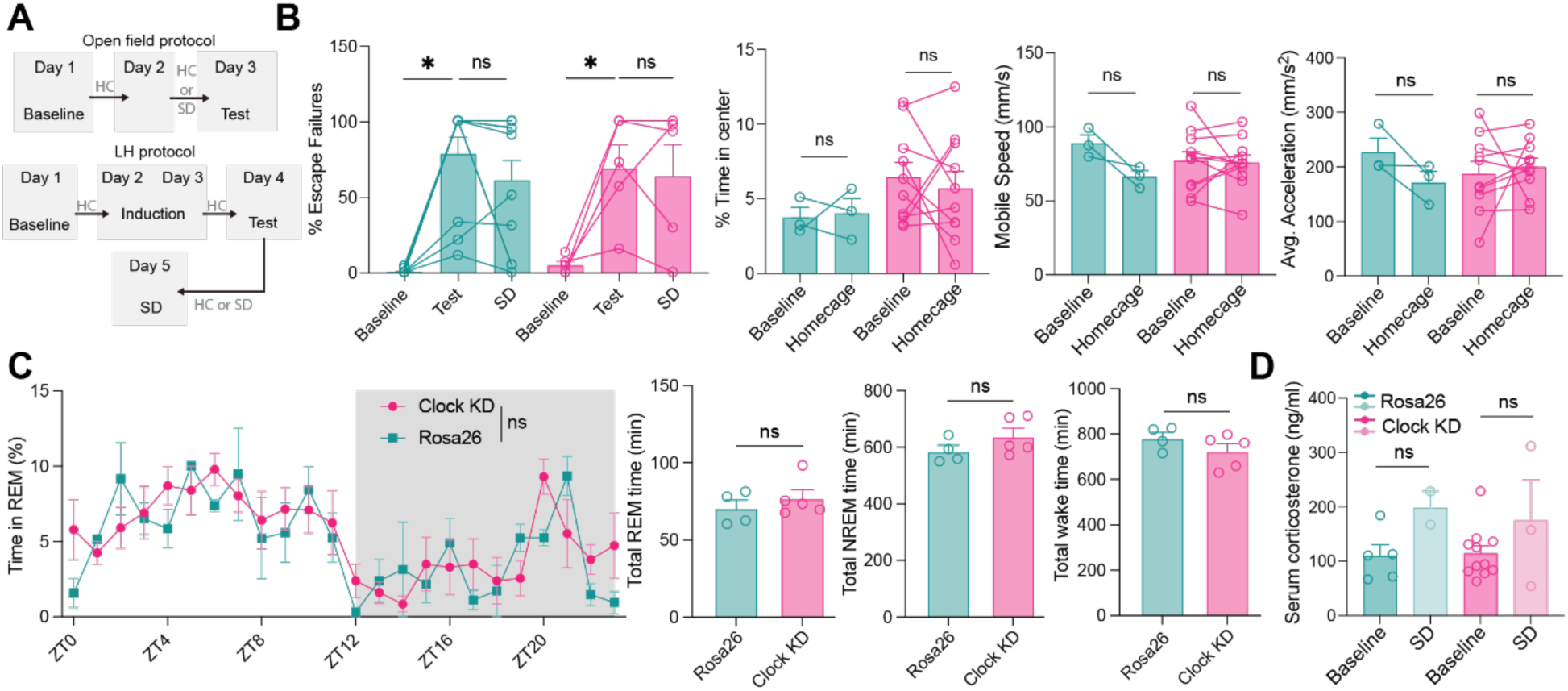
(**A**) Timeline of open field and learned helplessness behavior protocols. (**B**) Anxiety and locomotion are unchanged in controls (n=3) compared to Clock KD (n=11) mice when animals are allowed to sleep in their homecage. (**C**) Clock disruption in VTA dopaminergic neurons does not alter serum corticosterone levels. Serum corticosterone concentrations (ng/mL) measured at ZT12 in Rosa26 and Clock KD mice at baseline (n=5, n=11 respectively) and immediately following 12 hours of sleep deprivation (SD; n=2, n=3 respectively). (**D**) Percent and total time spent in REM state is unchanged in mice injected with Clock KD (n=5) compared to control (n=4) virus during adulthood. Total time spent in NREM, and wake state are also not significantly different between Clock KD mice and controls. ns, not significant, *p<0.05.

## Acknowledgements

This research was funded in whole by Breakthrough Discoveries for thriving with Bipolar Disorder (BD²; https://ror.org/00z5dw933) [Grant ID: DG230211]. For the purpose of open access, the author has applied a CC BY public copyright license to all Author Accepted Manuscripts arising from this submission.

CE is funded by the Novo Nordisk Foundation (NNF25OC0100934). OCG is funded by a K99 Fellowship from the National Institute on Aging (K99AG086609). LEF is supported by the Clayton Foundation for Research, W.M. Keck Foundation, BD2 Foundation, WoodNext Foundation, the NSF (award 24323797), and the Waggoner Center for Alcohol and Addiction Research.

## Competing Interests

LEF is a founder and officer of NumbCorp, a scientific advisory board member of Modulight.bio, and receives consulting income for work related to Parkinson’s disease, optogenetics, and cellular biologics.

## Methods

### Mice

All mice are on a C57BL/6J background. DAT-cre mice (B6.SJL-Slc6a3tm1.1 (cre)Bkmn/J) were acquired from Jackson Laboratories (Stock #006660). Mice were group-housed (max 5 mice/cage) on a twelve-hour light/dark cycle with *ad libitum* access to food and water. Experimental procedures were approved by The University of Texas at Austin, Northwestern University, and Stanford University IACUC committees and complied with the US National Institutes of Health Guide for the Care and Use of Laboratory Animals.

### Cells for *in vitro* test

NIH3T3 (ATCC, CRL-1658) and N2A cells (ATCC,CCL-131) were maintained at 37°C in an incubator with humidified 5% CO_2_. Cells were grown in DMEM media + Glutamax (GIBCO) supplemented with 10% FBS (Gibco) and 1% Penicillin-Streptomycin (Cytiva HyClone), with supplemented DMEM here on referred to as complete DMEM (cDMEM). Cells were enzymatically passaged by trypsinization with TrypLE Express (Gibco).

### Design of sgRNAs and cloning

Full genomic sequences for each target gene were obtained from the UCSC Genome Browser^57^. All annotated splice variants were aligned and overlapping coding regions common to all isoforms were selected for sgRNA targeting. Coding sequences were submitted to CRISPOR^38,39^ and CHOPCHOP^40,41^ to identify candidate sgRNAs and corresponding protospacer adjacent motif (PAM) sites. For Staphylococcus aureus Cas9 (SaCas9) targeting, sgRNAs of 21 bp in length with NNGRRT PAM sequences were selected. Final sgRNAs were chosen based on predicted on-target efficiency, likelihood of inducing frameshift mutations, and minimal off-target activity.

Each sgRNA was synthesized as a pair of complementary oligonucleotides (sense: 5′-CACCG-[sgRNA]-3′; antisense: 5′-AAAC-[reverse complement sgRNA]-C-3′). sgRNA sequences used in this study are listed in Table below. Oligos were resuspended to a final concentration of 100 µM. For annealing, 1 µL of each oligo (sense and antisense) was combined with 1 µL of T4 polynucleotide kinase (PNK, NEB), 1 µL of 10× T4 ligase buffer (NEB), and 6.5 µL of nuclease-free water. The reaction was incubated at 37°C for 30 min for phosphorylation, followed by denaturation at 95°C for 5 min and gradual cooling to room temperature to allow annealing.

Annealed oligos were ligated into a BsaI-digested AAV-CMV-DIO-SaCas9-sgRNA vector using T4 DNA ligase (NEB) at room temperature for 10 min. Subsequently, 2 µL of the ligation product was electroporated into NEB Stable competent *E. coli*. Transformed cells were plated on LB agar containing carbenicillin and incubated overnight. Colonies were selected and cultured in LB with carbenicillin, and plasmid DNA was isolated using the ZymoPURE Plasmid Miniprep Kit (D4210). Correct insertion of sgRNA sequences was confirmed by Sanger sequencing (Plasmidsaurus).

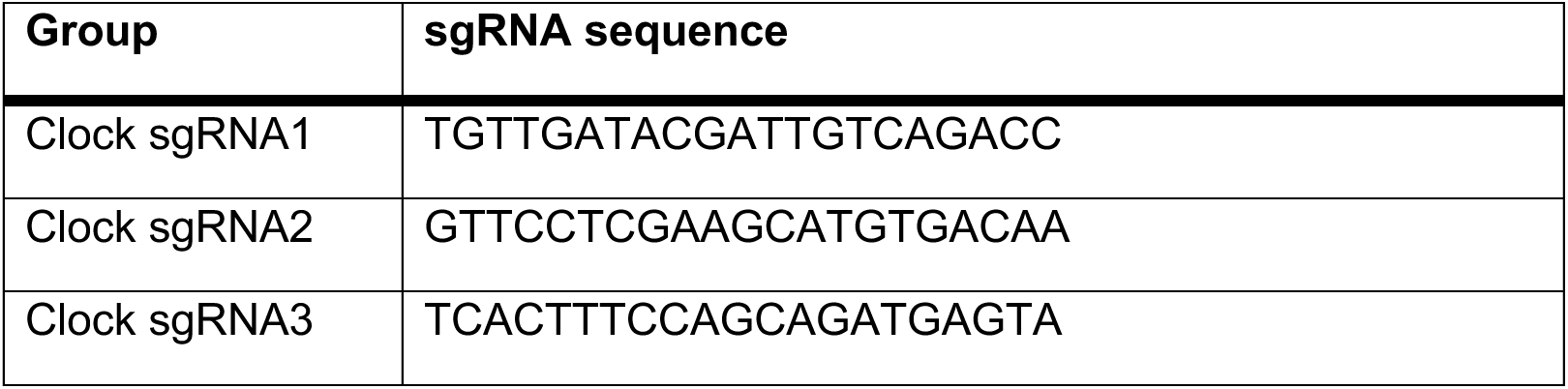

### Transfections

Lipofectamine LTX (Invitrogen) was used for all NIH3T3 and N2A cell transfections according to the manufacturer’s protocol. We plated 5×10^3^ cells/cm^2^ day before the transfection and used 1.2 µg of total DNA for 12-well plates. Cells were left to express endotoxin-free transfected DNA for 72 hrs before FACS sorting.

### Viral production

Most of the viruses used in editing efficiency validation were prepared in-house. Commercial preparations were contracted to Packgene and Vectorbuilder. All AAV produced with serotype DJ packaging vector with AAV2 ITRs.

In-house production was achieved using standard triple transfection of HEK293T/17 cells and iodixanol gradient ultracentrifuge purification. Briefly, HEK293T cells were grown on either 3 or 9 plates, 15 cm in diameter (Fisher Scientific, 12-600-004) until 70-80% confluency was reached. For 3 x 15 cm plates, 390 µl of polyethylenimine (PEI, Kyfora Bio, 260085) was used to transfect HEK293T cells. For AAV transfection, the following DNA mix was used: 50 µg of pAdDeltaF6 helper plasmid, 50 µg of pAAV-DJ Rep/Cap plasmid and 30 µg of payload plasmid. The DMEM and PEI were mixed in 1.5 ml of uncomplemented DMEM per 3 plates and left at RT for 10-20 mins before being added to 30 ml of pre-warmed cDMEM. 10 ml of this solution was then added to each plate and incubated for 72 hrs at 37°C in 5% CO_2_. Cells were then scraped off the plate with a cell lifter (Celltreat, MSPP-229306), collected in PBS and then pelleted at 800 x g for 10 mins. Cells were then combined and centrifuged again at 800 x g for 10 mins. If needed, cell pellets were frozen at -80°C as a pause point, before thawing at 37°C for 2 mins. Pellets were lysed in buffer containing 150 mM NaCl and 50 mM Tris-HCl and underwent two freeze-thaw cycles before being incubated with 50 U/ml benzonase nuclease (Sigma-Aldrich, E1014) for 30 mins at 37°C. Tubes were swirled by hand every 10 mins during incubation. Particles were cleared by centrifugation at 3000 x g for 15 mins, and clear supernatant was filtered through a 0.45 µm PES filter (VWR, 76479-020). The clarified supernatant was loaded on top of a discontinuous iodixanol gradient made up of 15%, 25%, 40% and 60% iodixanol layers in a 35 ml Quick-seal ultracentrifuge tube (Thermo Scientific, 03-989). This was spun at 350,000 x g for 90 mins at 12°C in a SorvallWX+ ultracentrifuge using maximum acceleration and deceleration. 2.5-3 ml of viral particles was collected by withdrawing the clear 40% fraction using an 18-gauge needle. Before purifying the recovered virus fraction through filtering, 15 ml Amicon Ultra Centrifugal Filters, 100-kDa molecular weight cut-off (Millipore, UFC9100), were incubated with 10 ml of 0.1% Pluronic F68 (MP Biomedicals, ICN2750049) in PBS for 10 mins at RT. This was then removed and replaced with 15 ml of 0.01% Pluronic F68 and spun at 3,000 x g for 5 min at 4°C. Flow-through was discarded, and 0.001% of Pluronic F68 with 200 mM NaCl was loaded and spun at 3,000 x g for 5 min at 4°C. The viral fraction was then loaded with 3-4 ml of 0.001% Pluronic F68 with 200 mM NaCl and centrifuged at 3,000 x g for 4 min at 4°C. Flow-through again was discarded, and the sample was centrifuged in 2 min intervals using previous conditions, until 500 µl of the sample remained. This retained sample was then filtered through a 0.5 ml Amicon Ultra Centrifugal Filter, 100-kDA molecular weight cut-off (Millipore, UFC510024) at 2,500 x g for 3 mins, and flow-through discarded. This was centrifuged in 2 min increments until the final volume of 70 µl was reached. The filter insert was then inverted and inserted into a fresh collection tube, which was centrifuged at 2,000 x g for 2 min. Purified virus was aliquoted into 2 µl aliquots and stored at - 80°C. Genomic titer was determined by ddPCR at PackGene Biotech, or by qPCR in house. Viral dilutions were calculated based on original titers calculated.

### Adult stereotaxic injections

Standard procedures were used to infuse AAV intracranially. Briefly, anesthesia induction and maintenance were with isoflurane. Buprenorphine, carprofen, and lidocaine were used for analgesia. 8-20 week old, anesthetized mice were placed in a stereotactic frame (Kopf Instruments or RWD). The skull was aligned using bregma and lambda. Craniotomies were performed using a hand drill. The following coordinates were used for stereotaxic viral injections targeting VTA (AP: -2.95, ML: ±0.5, DV: -4.5). Injections were made using a 10 µl Hamilton syringe with a 34-gauge beveled needle (World Precision Instruments). Unique syringes were dedicated to each viral mix. 1,000 nl of viral suspension was infused at a rate of 100 nl/min, with an additional needle dwell time of 10 min post-injection, before the needle is withdrawn. Skin was closed using nylon sutures. Standard, approved post-operative procedures were used.

### Nuclei isolation and FACS sorting

Midbrain tissue containing the VTA was rapidly dissected from mice, then isolated using a commercially available nuclei isolation kit (Miltenyi Biotec) according to the manufacturer’s instructions. Briefly, homogenized tissue was filtered through 70 μm and then 30 μm cell strainers to remove debris and large aggregates, followed by centrifugation to pellet nuclei. The resulting nuclei pellet was resuspended in a sorting buffer with magnetic beads and magnetic bound-nuclei were collected into a tube.

Fluorescence-activated cell sorting (FACS) was performed on a Sony SH800. Nuclei were first gated based on forward and back scatter parameters, and singlets were identified using FSC-H versus FSC-A gating. Reporter-positive nuclei were isolated by fluorescence-based sorting, with mCherry-positive cells selected for *in vitro* samples and GFP-positive nuclei selected for *in vivo* samples. GFP-positive nuclei were gated relative to fluorescence-negative controls and sorted into collection tubes containing PBS for downstream applications. Sorted samples were immediately processed for downstream assays, including genomic DNA amplification and sequencing.

### Amplicon generation and sequencing analysis

Whole genome amplification (WGA) was performed on the samples directly following FACS using the REPLI-g Advanced DNA Single Cell kit (QIAGEN) according to manufacturer’s instructions.

For generation of the specific amplicons, 1 µl of WGA DNA was diluted 1:50 and amplified (PCR 1) with Primestar Max Polymerase (Takara Bio) using the following protocol. Initial denaturation at 98°C for 30 sec, followed by 34 cycles of denaturation at 98°C for 10 sec, annealing at 65°C for 10 sec, and extension at 72°C for 10 sec. A final extension was performed at 72°C for 5 min. The product of PCR A was diluted 1:100 and amplified in a second PCR reaction, referred to as PCR B, using a nested PCR primers to decrease non-specific amplification and increase amplified DNA yield using the thermocycler protocol. The amplicons were gel extracted using the MinElute gel extraction kit (QIAGEN) and stored at -20°C or directly used in downstream processing. Primer sequences used for each target locus are listed below.

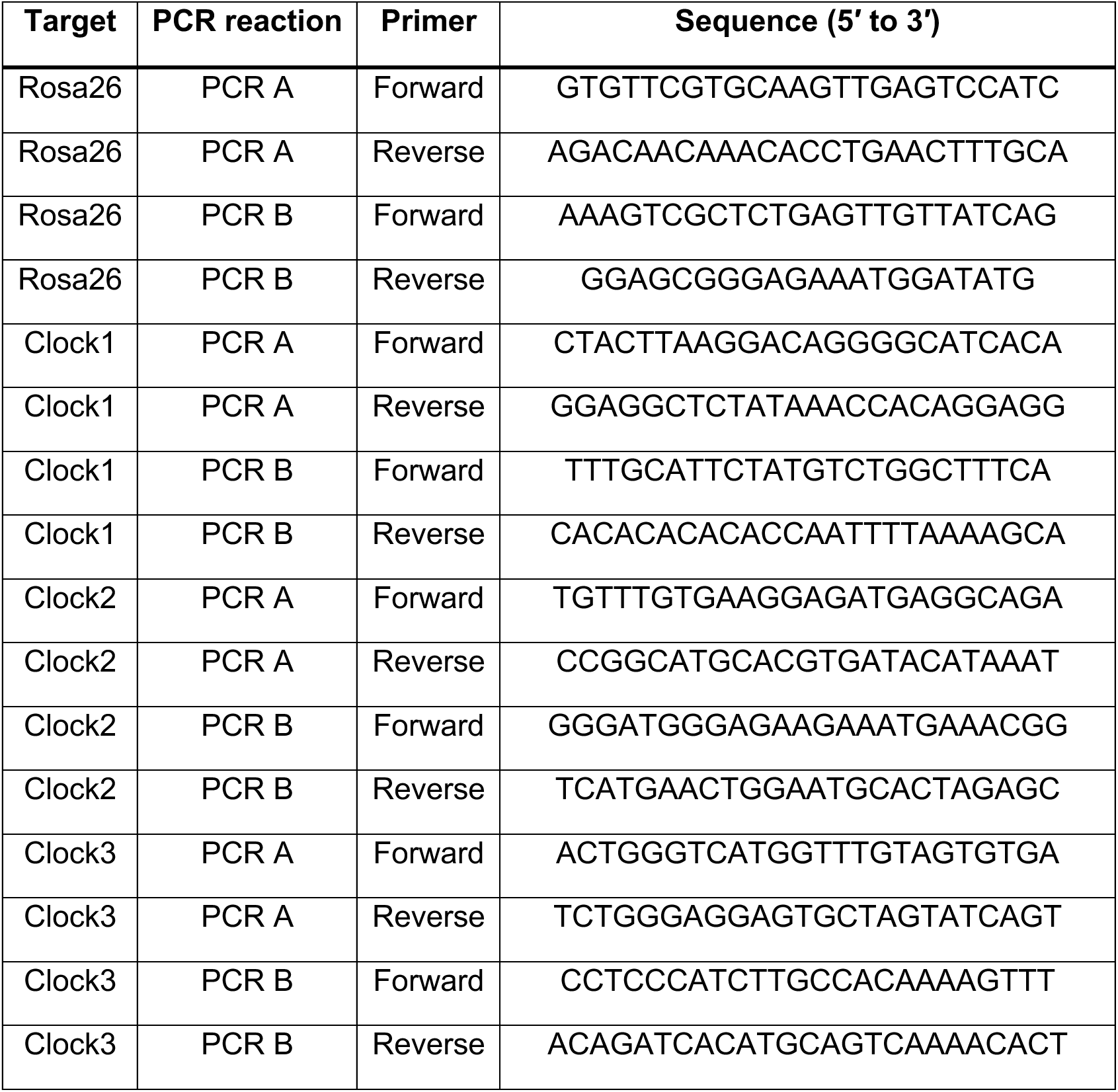

For sanger sequencing, purified PCR products were submitted to Eton Bioscience with either the forward or reverse primer, depending on sgRNA orientation, forward primer for sense sgRNAs and reverse primer for antisense sgRNAs. Sanger sequencing chromatograms were analyzed using TIDE, a sequence trace decomposition method^58^. Raw ab1 files obtained from Eton Bioscience were uploaded to the TIDE web tool, and indel frequencies were quantified relative to the corresponding wild-type control sample. For amplicon deep sequencing, purified PCR products were submitted to Plasmidsaurus for amplicon sequencing. Then, amplicon sequencing data were analyzed using CRISPRESSO^59^, which aligns sequencing reads to the expected amplicon reference sequence and quantifies genome editing outcomes at target site. The result output was used to determine the frequency and distribution of insertions and deletions at each target locus.

### Exome sequencing

Mice were injected with AAV delivering SaCas9 and 4 sgRNAs, ROSA and Clock1-3 and incubated 6 weeks to allow expression and editing to occur. Nuclei from injected mice were isolated using the Nuclei extraction and enrichment kit (Miltenyi Biotec). Nuclei underwent fluorescence-activated cell sorting to separate wildtype and SaCas9-sgRNA-expressing cells. Genomic DNA was isolated using the REPLI-g Single Cell Kit (Qiagen). Exome-captured libraries were constructed using Genewiz exome sequencing service. PCR enriched libraries underwent Illumina sequencing and exome sequence data were analyzed at Azenta Life Sciences. Raw BCL files were converted to FASTQ format using bcl2fastq v2.19. Raw reads were trimmed using Trimmomatic 0.39. Cleaned reads were aligned to the GRCm38/mm10 reference genome, sorted, and PCR duplicates were removed using Sentieon 202308. Single nucleotide variants and indels were called using Sentieon 202308 (DNAscope algorithm). We then excluded single nucleotide variations, indels found in at least two conditions, and naturally occurring common indels.

We identified potential off-target sites for each sgRNA target sequence using Cas-OFFinder^42,43^, searching the GRCm38/mm10 mouse reference genome from the Genome Reference Consortium for sequences with up to 7 mismatches to our target sequences and allowing for RNA and DNA bulges up to 2 bp. We then cross-checked with the exome sequencing results for indels occurring within 50 bp upstream or downstream from potential off-target sites.

### Immunohistochemistry

To validate knockout efficiency of different Clock sgRNAs, AAV-expressing mice were intra-peritoneally injected and deeply anesthetized with euthasol (Covetrus) before being transcardially perfused with 15 ml of ice-cold PBS followed by 15 ml of ice-cold 4% paraformaldehyde (PFA) solution in PBS. Brains were dissected out and post-fixed overnight in 4% PFA at 4°C before equilibration in sterile 30% sucrose in PBS for at least 24 hrs. Tissue was sectioned at 30 µm on a cryostat (Leica) and either stained and mounted with VECTASHIELD Antifade Mounting Medium with DAPI (Vector Laboratories), or stored in cryoprotectant (25% glycerol (Fisher Scientific), 30% ethylene glycol (Macron Fine Chemicals) in PBS, pH 6.7 and 0.2 µM filter sterilized). 10x and 60x fluorescence images were obtained using a Nikon ECLIPSE Ti2 confocal microscope. For each experiment, imaging settings were kept constant.

Slices undergoing immunohistochemistry were washed in PBS as free-floating sections before being blocked and permeabilized in blocking solution (0.3% Triton X-100 (Sigma-Aldrich) and 4% donkey serum (Jackson ImmunoResearch) in 1x PBS) at RT for 1 hour. Slices were then incubated overnight at 4°C in blocking solution with primary antibody. Sections were next washed in PBS and then incubated in blocking serum with the appropriate secondary antibody at room temperature for 90 min. Slices were washed 3x with PBS before being mounted and coverslipped using VECTASHIELD Antifade Mounting Medium with DAPI (Vector Laboratories). Primary antibodies used: rabbit anti-Clock (Genewiz, GTX636283, 1:150), mouse anti-HA tag (Invitrogen, 26183, 1:500). Secondary antibodies (all 1:500 dilution): Rhodamine Red-X donkey anti-rabbit (Jackson ImmunoResearch, 711-295-152), and Alexa Fluor 647 donkey anti-mouse (Jackson ImmunoResearch, 715-605-150).

### Neonatal stereotaxic injections and validation

Neonatal viral transduction was carried out in order to minimize invasiveness, increase surgical efficiency, and to more closely mimic the developmental timelines in humans carrying BD risk alleles. Standard procedures we have previously used were followed^44,45,60^. P1-3 mice were cryo-anesthetized and placed on a cooling pad. The virus was delivered at 100 nl/min, for 500 nl per side, using an UltraMicroPump (World Precision Instruments, Sarasota FL). The ventral tegmental area (VTA) was targeted in neonates by placing the needle approximately 0.1 mm anterior to lambda, ± 0.1 mm laterally from the midline, and 3.6 mm ventral to the skin surface. Coordinates were slightly adjusted based on age and size of each pup. Following the procedure, pups were warmed on a heating pad and returned to their home cages.

To validate successful neonatal VTA targeting in behavior animals, after all behavior tests were completed, mice were deeply anesthetized with isoflurane and transcardially perfused with 4% paraformaldehyde (PFA) in 0.1 M phosphate-buffered saline (PBS). Brains were post-fixed for 1-3 days in 4% PFA at 4°C and washed in PBS prior to sectioning at 60 μm on a vibratome (Leica Biosystems). Mice were euthanized all between 1-4 hours into the dark phase.

Sections were then washed in PBS as free-floating sections before being permeabilized with 0.2% Triton X-100 for 2 hours at RT and then blocked in 10% bovine serum albumin (BSA, Sigma-Aldrich), in PBS with 0.1% Triton X-100 for 2 hours at RT. Sections were then incubated for 16 hours overnight at 4°C in primary antibody solution with 5% BSA and 0.2% Triton X-100. The following day, tissues were rinsed in PBS, reacted with secondary antibodies for 2 hours at RT, rinsed again, and then mounted onto Superfrost Plus slides (ThermoFisher Scientific). Sections were dried and coverslipped under glycerol:TBS (9:1) with Hoechst 33342 (2.5 μg/ml, ThermoFisher Scientific). Primary antibodies used: goat anti-RFP (Rockland, 200-101-379, 1:500), sheep anti-tyrosine hydroxylase (Abcam, ab113, 1:1000). Secondary antibodies used (all 1:1000 dilution): Alexa Fluor 594 donkey anti-goat (Invitrogen, A-11058), Alexa Fluor 488 donkey anti-sheep (Invitrogen, A-11015).

Post hoc analysis was carried out for all animals that underwent behavior testing. Mice with fewer than an average of 30 tdTomato+ ROIs per section were omitted from the behavior tests (**Supplementary Fig. 2C-D**). Out of 11 injected control mice and 26 injected Clock KD mice, we excluded 2 and 5 mice respectively due to suboptimal expression of the fluorophore AAV. No other animals were excluded from the analyses.

### Behavioral assays

All behavior tests were done at the beginning of the active phase of the circadian cycle (ZT12). Mice were placed in reverse light dark cycle at least 1 week before testing began. To assess locomotor and anxiety-like behaviors, mice (>P60) were placed in an open field arena under infrared illumination at the beginning of the active phase of the circadian cycle. Mice explored the arena for 20 minutes with video recording using a raspberry pi camera (50 fps). Prior to testing periods, 2 habituation periods were done 2 days before testing day. For habituation, mice spent a minimum of 1 hr in their homecage in the behavior room and explored the open field arena for 10 minutes on each day.

To assess the baseline propensity to avoid unpleasant stimuli (foot shocks), mice were placed in an active-passive avoidance shuttle box (MazeEngineers) and 30-45 escapable foot shocks (0.3 mA, 10 sec, random 5-20 sec inter-stimulus intervals) were delivered. Escapes were scored when the animal shuttled between compartments during the stimulus time window. The stimulus automatically terminated when the mice shuttled to the other compartment. Failures were scored when the mice failed to shuttle to the other compartment in the duration of the foot shock. Following a baseline testing session, two induction sessions (1 session per day; 360 inescapable foot shocks per session; 0.3 mA, 3 sec; random 3-15 sec inter-stimulus intervals) were conducted. A day after induction, a repeat test session with avoidable stimuli was conducted. All test sessions were completed at the beginning of the active phase of the circadian cycle.

### Sleep deprivation paradigm

Custom-built sleep deprivation devices were built as previously described^45^. In brief, each device consists of a cylinder (30.5 cm height, 19.7 cm inner diameter) and a circular plate (17.8 cm diameter) acting as the base for the motor and rotating beam. A DC motor (12 V) was attached to the base, and a plastic bar (0.5 cm x 0.5 cm x 17.5 cm) served as the rotating beam. A platform (3 cm diameter) was placed 7 cm above the base and within reach of the wire rack holding food and water. Mice were free to access food and drink in the duration of time in the chamber. On the day of the acute sleep deprivation, mice were placed into the chamber from light onset to offset (ZT 0-12) in a room with standard illumination and temperature control. At the end of the sleep deprivation period, the lights were switched to infrared for further behavioral evaluation. All mice were returned to their home cage for 5-10 minutes before behavior testing was done.

### EEG/EMG surgery and recording

Neonatal injections were performed as described above. At 6 weeks of age, EEG/EMG surgeries were performed. The mice were anesthetized with isoflurane (induction 4%; 4L/min O2, maintenance 1.5-2.5; 2L/min O2) and received a subcutaneous injection of Buprenorphine Slow Release (0.5 mg/kg) before being placed in the stereotaxic rig (David Kopf Instruments, Tujunga, CA) on a heating pad. Antibiotic ointments were applied, and 0.3 mL of saline was injected subcutaneously to prevent dehydration. The headsets, consisting of stainless steel mini-screws (US Micro Screw) and metal wires (316SS/44T, Medwire) for EEG and EMG, were implanted on the skull above the frontal (AP + 1.5 mm; ML 1.5 mm) and temporal (AP - 2 mm; ML 2 mm) lobes. The metal wires were placed intramuscularly bilaterally in the nuchal muscles for EMG. The EEG/EMG headset was mounted on the skull with Metabond (Parkell) and Fusion Flo (Prevest DenPro). After surgery, the mice were placed in a separate cage on a heating pad for recovery and subsequently housed with littermates.

### EEG/EMG recordings and analyses

Two weeks after surgery, the mice were single housed in the EEG/EMG recording cages for 3 days prior to the recording for habituation. EEG/EMG signals were recorded for 24 hours to include both light and dark cycles. EEG/EMG signals were amplified using a multi-channel amplifier (Grass Instruments) and recorded with VitalRecorder (Kissei Comtec Co.) at a sampling rate of 256 Hz. Raw EEG/EMG data were converted to SleepSign (Kissei Comtec Co.), exported to MATLAB (MathWorks, Natick, MA, USA), and preprocessed. The EEG and EMG recordings were digitally band-pass filtered (EEG: 1-126 Hz; EMG: 10-40 Hz) using a second-order Butterworth filter. Further, zero-phase filtering was performed with forward and reverse application of the filter coefficients. Sleep recordings were scored using AccuSleep^61^. A minimum of 1 hour was scored manually in 4-second epochs (including wake, NREM, and REM sleep) according to standard criteria^62^. These data were used as labeled input to train the AccuSleep model for state classification. The remainder of the recording was scored automatically and manually checked to ensure appropriate scoring. Sleep-state duration metrics were computed and visualized in Python using Visual Studio Code.

### Acute brain slice preparation

Male and female neonatal mice were intracranially injected as described above with a mix of AAV-CAG-DIO-tdTomato and AAV-DIO-SaCas-Rosa26-sgRNA (Rosa26 controls) or AAV-CAG-DIO-tdTomato and AAV-DIO-SaCas-Clock-sgRNA 3 (Clock KD). At 8-10 weeks of age, the mice were deeply anesthetized for acute midbrain slice preparation using ketamine (75 mg/kg) and dexmedetomidine (0.25 mg/kg). Cardiac blood collection was then performed from the right ventricle to collect serum for the corticosterone ELISA assay, and then mice were quickly perfused with an ice-cold, oxygenated choline-based solution containing (in mM): 110 choline chloride, 25 NaHCO_3_, 2.5 KCl, 1.25 NaH_2_PO_4_, 0.5 CaCl_2_, 7 MgCl_2_, 11.6 sodium ascorbate, 3.1 sodium pyruvate, and 25 glucose, saturated with 95%O_2_/5% CO_2_ (pH 7.4; 290-300 mOsm). Following perfusion, brains were dissected and horizontal slices (220 µm) were prepared using a Leica VT 1200S vibratome in the choline solution. Slices recovered at 34°C for 1 hour in oxygenated artificial cerebrospinal fluid (ACSF in mM) containing: 126 NaCl, 21.4 NaHCO_3_, 2.5 KCl, 1.2 NaH_2_PO_4_, 2.4 CaCl_2_, 1.2 MgSO_4_ and 11.1 glucose, saturated with 95% O_2_/5% CO_2_ (pH 7.4; 290-300 mOsm), and then were held submerged at RT until being transferred to the recording chamber.

### Electrophysiological recordings

Horizontal midbrain slices were continuously perfused (1.5-2 mL/min) with oxygenated artificial cerebrospinal fluid (ACSF) maintained at 29-30°C. Patch pipettes (2–5 MΩ) were filled with an internal solution containing (in mM): 117 K-gluconate, 2.8 NaCl, 5 MgCl₂, 0.2 CaCl₂, 2 Na₂-ATP, 0.3 Na-GTP, 0.6 EGTA, and 10 HEPES (pH 7.25–7.28; 265–280 mOsm). Whole-cell patch-clamp recordings were obtained from tdTomato-labeled neurons in the ventral tegmental area (VTA). Neurons were recorded in either voltage-clamp or current-clamp configuration to assess intrinsic membrane properties and excitability. In voltage-clamp recording, dopamine (DA) neurons were held at -70 mV to minimize spontaneous firing. After breaking in to whole cell mode, intrinsic properties were assessed in voltage-clamp: hyperpolarization-activated inward current (H-current) was evoked using a 750 ms hyperpolarizing step from -40 mV to -120 mV; currents underlying the afterhyperpolarization (AHP) were measured using a 100 ms depolarizing step from -50 mV to 0 mV; input resistance and series resistance were measured using a -5 mV hyperpolarizing step from -70 mV (100 ms duration). In current-clamp recordings, resting membrane potential (RMP) was measured after establishing whole-cell configuration, the calculated liquid junction potential at 30°C was 13 mV; membrane potentials were not corrected for junction potential. Rheobase (current required to elicit the first action potential) and action potential (AP) threshold were determined using a depolarizing current ramp (+450 pA over 1 s), applied from resting membrane potential. Evoked firing was assessed using 500 ms current steps ranging from -100 to +600 pA, in 50 pA increments.

### Serum corticosterone ELISA

Serum corticosterone concentrations were quantified using the DetectX® Corticosterone Multi-Format ELISA Kit (Arbor Assays, K014-H1). Samples were treated with Dissociation Reagent and diluted 1:100 in 1X Assay Buffer. All samples and standards were run in duplicate using the 50 μL format. Absorbance was measured at 450 nm and concentrations were calculated from a nine-point standard curve using four-parameter logistic (4PL) curve fitting in GraphPad Prism. Blood was collected at ZT12; samples from sleep-deprived animals were collected immediately following the 12-hour sleep deprivation period.

### Quantification and statistical analyses

#### Quantification of behavior

For evaluating locomotor behaviors, Toxtrac^63,64^ was used to track the animal’s position, defined by its body center, and quantify the distance traveled in each session. Custom MATLAB scripts were used to quantify the time spent in the center of the open field, defined as the middle third of the box. Percent of escape failures were quantified as the number of escape failures divided by the sum of escape failures and successful escapes. This was calculated for baseline and post-induction testing sessions.

#### Statistical analyses

Animals or cells were randomly assigned to treatment groups. Group statistical analyses were done using GraphPad Prism 9 software (GraphPad, Lajolla, CA). All data are expressed as mean +/- SEM or individual plots. For two-group comparisons, statistical significance was determined by two-tailed Student’s t-tests. For multiple group comparisons, one-way or two-way analysis of variance (ANOVA) tests were used for normally distributed data, followed by post hoc analyses. For non-normally distributed data, non-parametric tests for the appropriate group numbers were used, as noted in the legends. p<0.05 was considered statistically significant.

## Notes

### Competing Interest Statement

The authors have declared no competing interest.

### Summary of Updates

The details of the sponsorship and author affiliations have been updated. The licensing for the preprint has been updated.

## References

1. Tonon, A. C. et al. Sleep and circadian disruption in bipolar disorders: From psychopathology to digital phenotyping in clinical practice. Psychiatry Clin Neurosci 78, 654–666 (2024).

2. Takaesu, Y. Circadian rhythm in bipolar disorder: A review of the literature. Psychiatry and Clinical Neurosciences 72, 673–682 (2018).

3. Gonzalez, R., Gonzalez, S. D. & McCarthy, M. J. Using Chronobiological Phenotypes to Address Heterogeneity in Bipolar Disorder. Molecular Neuropsychiatry 5, 72–84 (2020).

4. Harvey, A. G. Sleep and Circadian Rhythms in Bipolar Disorder: Seeking Synchrony, Harmony, and Regulation. American Journal of Psychiatry 165, 820–829 (2008).

5. Jones, S. H., Hare, D. J. & Evershed, K. Actigraphic assessment of circadian activity and sleep patterns in bipolar disorder. Bipolar Disorders 7, 176–186 (2005).

6. Salvatore, P. et al. Circadian activity rhythm abnormalities in ill and recovered bipolar I disorder patients. Bipolar Disorders 10, 256–265 (2008).

7. Mansour, H. A. et al. Circadian Phase Variation in Bipolar I Disorder. Chronobiology International 22, 571–584 (2005).

8. Boudebesse, C. et al. Correlations between objective and subjective sleep and circadian markers in remitted patients with bipolar disorder. Chronobiology International 31, 698–704 (2014).

9. Song, Y. M. et al. Causal dynamics of sleep, circadian rhythm, and mood symptoms in patients with major depression and bipolar disorder: insights from longitudinal wearable device data. eBioMedicine 103, 105094 (2024).

10. Harvey, A. G. Sleep and Circadian Functioning: Critical Mechanisms in the Mood Disorders? Annual Review of Clinical Psychology 7, 297–319 (2011).

11. Bauer, M. et al. Temporal relation between sleep and mood in patients with bipolar disorder. Bipolar Disorders 8, 160–167 (2006).

12. Murray, G. & Harvey, A. Circadian rhythms and sleep in bipolar disorder. Bipolar Disorders 12, 459–472 (2010).

13. Kim, Y. H. & Lazar, M. A. Transcriptional Control of Circadian Rhythms and Metabolism: A Matter of Time and Space. Endocr Rev 41, 707–732 (2020).

14. Baser, K. H. C., Haskologlu, I. C. & Erdag, E. Molecular Links Between Circadian Rhythm Disruption, Melatonin, and Neurodegenerative Diseases: An Updated Review. Molecules 30, 1888 (2025).

15. Lee, Y., Chen, R., Lee, H. & Lee, C. Stoichiometric Relationship among Clock Proteins Determines Robustness of Circadian Rhythms *. Journal of Biological Chemistry 286, 7033–7042 (2011).

16. Fagiani, F. et al. Molecular regulations of circadian rhythm and implications for physiology and diseases. Sig Transduct Target Ther 7, 41 (2022).

17. Gekakis, N. et al. Role of the CLOCK Protein in the Mammalian Circadian Mechanism. Science 280, 1564–1569 (1998).

18. McClung, C. A. Circadian genes, rhythms and the biology of mood disorders. Pharmacology & Therapeutics 114, 222–232 (2007).

19. Waddington Lamont, E., Legault-Coutu, D., Cermakian, N. & Boivin, D. B. The role of circadian clock genes in mental disorders. Dialogues Clin Neurosci 9, 333–342 (2007).

20. Roybal, K. et al. Mania-like behavior induced by disruption of CLOCK. Proc. Natl. Acad. Sci. U.S.A. 104, 6406–6411 (2007).

21. Spencer, S. et al. A mutation in CLOCK leads to altered dopamine receptor function. Journal of Neurochemistry 123, 124–134 (2012).

22. Van Enkhuizen, J., Milienne-Petiot, M., Geyer, M. A. & Young, J. W. Modeling bipolar disorder in mice by increasing acetylcholine or dopamine: chronic lithium treats most, but not all features. Psychopharmacology 232, 3455–3467 (2015).

23. Charrier, A., Olliac, B., Roubertoux, P. & Tordjman, S. Clock Genes and Altered Sleep–Wake Rhythms: Their Role in the Development of Psychiatric Disorders. International Journal of Molecular Sciences 18, 938 (2017).

24. Mukherjee, S. et al. Knockdown of Clock in the Ventral Tegmental Area Through RNA Interference Results in a Mixed State of Mania and Depression-Like Behavior. Biological Psychiatry 68, 503–511 (2010).

25. Nair-Roberts, R. G. et al. Stereological estimates of dopaminergic, GABAergic and glutamatergic neurons in the ventral tegmental area, substantia nigra and retrorubral field in the rat. Neuroscience 152, 1024–1031 (2008).

26. Walsh, J. J. & Han, M. H. THE HETEROGENEITY OF VENTRAL TEGMENTAL AREA NEURONS: PROJECTION FUNCTIONS IN A MOOD-RELATED CONTEXT. Neuroscience 282, 101–108 (2014).

27. Atasoy, D., Aponte, Y., Su, H. H. & Sternson, S. M. A FLEX Switch Targets Channelrhodopsin-2 to Multiple Cell Types for Imaging and Long-Range Circuit Mapping. J Neurosci 28, 7025–7030 (2008).

28. Schnütgen, F. et al. A directional strategy for monitoring Cre-mediated recombination at the cellular level in the mouse. Nat Biotechnol 21, 562–565 (2003).

29. Hunker, A. C. et al. Conditional Single Vector CRISPR/SaCas9 Viruses for Efficient Mutagenesis in the Adult Mouse Nervous System. Cell Rep 30, 4303–4316.e6 (2020).

30. Grieger, J. C. & Samulski, R. J. Packaging Capacity of Adeno-Associated Virus Serotypes: Impact of Larger Genomes on Infectivity and Postentry Steps. J Virol 79, 9933–9944 (2005).

31. Lusby, E., Fife, K. H. & Berns, K. I. Nucleotide sequence of the inverted terminal repetition in adeno-associated virus DNA. J Virol 34, 402–409 (1980).

32. Friedrich, G. & Soriano, P. Promoter traps in embryonic stem cells: a genetic screen to identify and mutate developmental genes in mice. Genes Dev. 5, 1513–1523 (1991).

33. Soriano, P. Generalized lacZ expression with the ROSA26 Cre reporter strain. Nat Genet 21, 70–71 (1999).

34. Ran, F. A. et al. In vivo genome editing using Staphylococcus aureus Cas9. Nature 520, 186–191 (2015).

35. Shi, J. et al. Clock genes may influence bipolar disorder susceptibility and dysfunctional circadian rhythm. American J of Med Genetics Pt B **147B**, 1047–1055 (2008).

36. McCarthy, M. J., Nievergelt, C. M., Kelsoe, J. R. & Welsh, D. K. A Survey of Genomic Studies Supports Association of Circadian Clock Genes with Bipolar Disorder Spectrum Illnesses and Lithium Response. PLoS ONE 7, e32091 (2012).

37. Bult, C. J., Blake, J. A., Smith, C. L., Kadin, J. A. & Richardson, J. E. Mouse Genome Database (MGD) 2019. Nucleic Acids Res 47, D801–D806 (2019).

38. Concordet, J.-P. & Haeussler, M. CRISPOR: intuitive guide selection for CRISPR/Cas9 genome editing experiments and screens. Nucleic Acids Res 46, W242–W245 (2018).

39. Haeussler, M. et al. Evaluation of off-target and on-target scoring algorithms and integration into the guide RNA selection tool CRISPOR. Genome Biol 17, 148 (2016).

40. Labun, K. et al. CHOPCHOP v3: expanding the CRISPR web toolbox beyond genome editing. Nucleic Acids Research 47, W171–W174 (2019).

41. Montague, T. G., Cruz, J. M., Gagnon, J. A., Church, G. M. & Valen, E. CHOPCHOP: a CRISPR/Cas9 and TALEN web tool for genome editing. Nucleic Acids Research 42, W401–W407 (2014).

42. Cho, S. W. et al. Analysis of off-target effects of CRISPR/Cas-derived RNA-guided endonucleases and nickases. Genome Res. 24, 132–141 (2014).

43. Bae, S., Park, J. & Kim, J.-S. Cas-OFFinder: a fast and versatile algorithm that searches for potential off-target sites of Cas9 RNA-guided endonucleases. Bioinformatics 30, 1473–1475 (2014).

44. Wu, M. et al. Attenuated dopamine signaling after aversive learning is restored by ketamine to rescue escape actions. eLife 10, e64041 (2021).

45. Wu, M. et al. Dopamine pathways mediating affective state transitions after sleep loss. Neuron S0896627323007584 (2023) doi:10.1016/j.neuron.2023.10.002.

46. Milo, T. et al. Longitudinal hair cortisol in bipolar disorder and a mechanism based on HPA dynamics. iScience 27, 109234 (2024).

47. Eban-Rothschild, A., Rothschild, G., Giardino, W. J., Jones, J. R. & De Lecea, L. VTA dopaminergic neurons regulate ethologically relevant sleep–wake behaviors. Nat Neurosci 19, 1356–1366 (2016).

48. Morales, M. & Margolis, E. B. Ventral tegmental area: cellular heterogeneity, connectivity and behaviour. Nat Rev Neurosci 18, 73–85 (2017).

49. Perugi, G. et al. Mixed Features in Patients With a Major Depressive Episode: the BRIDGE-II-MIX Study. J. Clin. Psychiatry 76, e351–e358 (2015).

50. Vázquez, G. H. et al. Mixed symptoms in major depressive and bipolar disorders: A systematic review. Journal of Affective Disorders 225, 756–760 (2018).

51. Sulaman, B. A., Wang, S., Tyan, J. & Eban-Rothschild, A. Neuro-orchestration of sleep and wakefulness. Nat Neurosci 26, 196–212 (2023).

52. Walker, W. H., Walton, J. C., DeVries, A. C. & Nelson, R. J. Circadian rhythm disruption and mental health. Transl Psychiatry 10, 28 (2020).

53. Mullins, N. et al. Genome-wide association study of more than 40,000 bipolar disorder cases provides new insights into the underlying biology. Nat Genet 53, 817–829 (2021).

54. Craddock, N. & Sklar, P. Genetics of bipolar disorder. The Lancet 381, 1654–1662 (2013).

55. Watson, C. J. et al. Phenomics-based quantification of CRISPR-induced mosaicism in zebrafish. Cell Syst 10, 275–286.e5 (2020).

56. Mehravar, M., Shirazi, A., Nazari, M. & Banan, M. Mosaicism in CRISPR/Cas9-mediated genome editing. Developmental Biology 445, 156–162 (2019).

57. Casper, J. et al. The UCSC Genome Browser database: 2026 update. Nucleic Acids Research 54, D1331–D1335 (2026).

58. Brinkman, E. K. & Van Steensel, B. Rapid Quantitative Evaluation of CRISPR Genome Editing by TIDE and TIDER. in CRISPR Gene Editing (ed. Luo, Y.) vol. 1961 29–44 (Springer New York, New York, NY, 2019).

59. Clement, K. et al. CRISPResso2 provides accurate and rapid genome editing sequence analysis. Nat Biotechnol 37, 224–226 (2019).

60. Wu, M., Minkowicz, S., Dumrongprechachan, V., Hamilton, P. & Kozorovitskiy, Y. Ketamine Rapidly Enhances Glutamate-Evoked Dendritic Spinogenesis in Medial Prefrontal Cortex Through Dopaminergic Mechanisms. Biological Psychiatry 89, 1096–1105 (2021).

61. Barger, Z., Frye, C. G., Liu, D., Dan, Y. & Bouchard, K. E. Robust, automated sleep scoring by a compact neural network with distributional shift correction. PLoS ONE 14, e0224642 (2019).

62. Mang, G. M. & Franken, P. Sleep and EEG Phenotyping in Mice. CP Mouse Biology 2, 55–74 (2012).

63. Rodriguez, A., Zhang, H., Klaminder, J., Brodin, T. & Andersson, M. ToxId: an efficient algorithm to solve occlusions when tracking multiple animals. Sci Rep 7, 14774 (2017).

64. Rodriguez, A., et al. *ToxTrac* : A fast and robust software for tracking organisms. Methods Ecol Evol 9, 460–464 (2018).

